# Chemi-Northern: a versatile chemiluminescent northern blot method for analysis and quantitation of RNA molecules

**DOI:** 10.1101/2023.10.10.561763

**Authors:** Katherine M. McKenney, Robert P. Connacher, Elise B. Dunshee, Aaron C. Goldstrohm

## Abstract

This report describes a chemiluminescence-based detection method for RNAs on northern blots, designated Chemi-Northern. This approach builds on the simplicity and versatility of northern blotting, while dispensing of the need for expensive and cumbersome radioactivity. RNAs are first separated on denaturing gel electrophoresis, transferred to a nylon membrane, and then hybridized to a biotinylated RNA or DNA antisense probe. Streptavidin conjugated with horseradish peroxidase and enhanced chemiluminescence substrate are then used to detect the probe bound to the target RNA. Our results demonstrate the versatility of this method in detecting natural and engineered RNAs expressed in cells, including messenger and noncoding RNAs. We show that Chemi-Northern detection is sensitive and fast, detecting attomole amounts of RNA in as little as 1 second, with high signal intensity and low background. The dynamic response displays excellent linearity. Using Chemi-Northern, we measure the significant, reproducible reduction of mRNA levels by human sequence-specific RNA-binding proteins, PUM1 and PUM2. Additionally, we measure the interaction of endogenous poly(A) binding protein, PABPC1, with poly-adenylated mRNA. Thus, the Chemi-Northern method provides a versatile, simple, cost-effective method to enable researchers to detect and measure changes in RNA expression, processing, binding, and decay of RNAs.

## Introduction

The need to detect and quantitate RNA molecules is pervasive in biological and medical research. Fortunately, multiple approaches are available to researchers. Each method has inherent strengths and limitations that an investigator should consider. For example, reverse transcription coupled with quantitative polymerase chain reaction (RT-qPCR) and whole transcriptome RNA sequencing (RNA-Seq) are powerful tools (1,2); however, those methods require expensive equipment and reagents and careful optimization of amplification and library generation steps. Moreover, RT-qPCR and RNA-Seq detect small amplicons, limiting their ability to distinguish overlapping RNA isoforms, RNA processing and decay intermediates, and post-transcriptional modifications. Long-read sequencing technologies hold promise, though their high cost, technical, and equipment requirements remain a barrier to widespread routine application (1).

A facile approach for RNA analysis that can be widely implemented would be beneficial to researchers. Ideally the method should be versatile in its ability to detect diverse RNA species from a variety of sources. It should offer sensitive detection and reproducible quantitation of RNA levels. Utilization of such a method would be improved by low cost, simplicity, and wide availability of reagents and equipment. For more than 40 years, the stalwart method that fits these criteria is northern blotting (3,4). Northern blotting can be used to measure the size and level of intact RNAs, along with changes in their processing, modification, and metabolism. The procedure is simple (**Figure 1**). First, an antisense nucleic acid probe corresponding to the target RNA is made. Second, the RNA is purified from the sample. Third, the RNA is physically separated based on size (i.e. charge to length ratio) via denaturing gel electrophoresis. Fourth, the RNA is transferred from the gel to a positively charged membrane. Fifth, the RNA of interest is detected through hybridization to an antisense probe coupled with a detectable signal.

**Figure 1.**
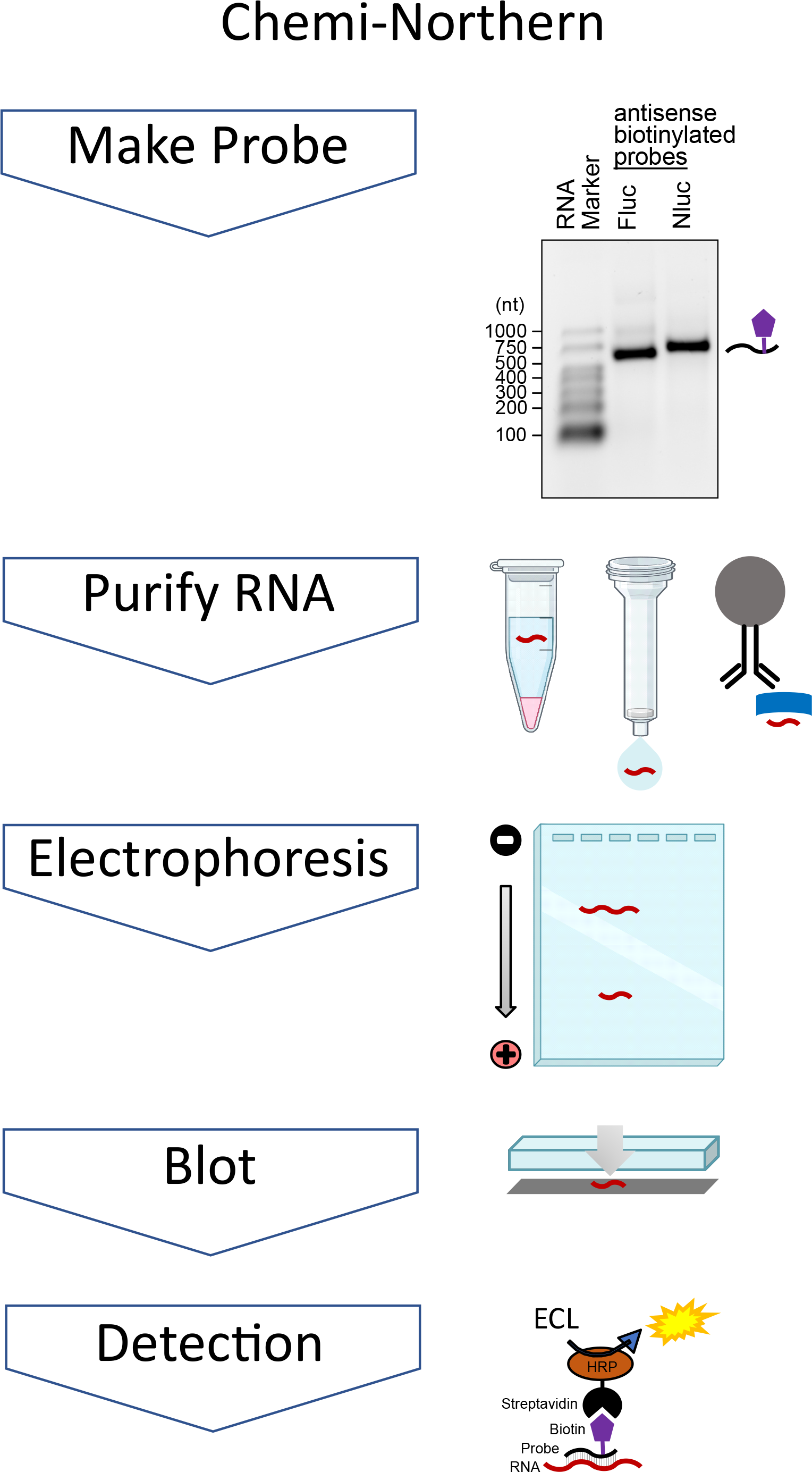
Overview of RNA detection by Chemi-Northern method. The Chemi-Northern protocol consists of 5 steps: 1) Generation of biotinylated (purple pentagon) antisense nucleic acid probe (black line). As an example, an ethidium bromide stained denaturing formaldehyde-MOPS agarose gel of the firefly luciferase (Fluc) and nano-luciferase (Nluc) probes used in this study are shown on the right. An RNA size marker is included on the left, with sizes indicated in nucleotides (nt). 2) Extraction and purification of RNA (red line). As shown on the right, RNAs can be purified from a diverse sources using a variety of methods including organic phase separation (e.g. trizol reagent), spin column, or bead based methods (e.g. RNA coimmunoprecipitation assays). 3) Separation of RNA by charge to length ratio by denaturing formaldehyde-MOPS agarose gel electrophoresis. 4) Blotting transfer and immobilization of RNA to positively charged nylon membrane. 5) Visualization of target RNA hybridized to the biotinylated probe using streptavidin-HRP conjugate and enhanced chemiluminescence (ECL) detection. Gel, column, and test tube icons were created by Biorender.com under academic license agreement NE25MGJNLD.

Traditionally, probes for northern blotting were labeled by incorporation of radioactive ^32^P, which enabled detection by autoradiography or, subsequently, by phosphorimaging (3,4). However, several factors have diminished the adoption of radioactive northern blots. First, use of radioactive material requires special licensing, regulatory oversight, procedures, equipment, and facilities. Second, the cost of radioactive nucleotides has substantially increased. Third, the short 14 day half-life of ^32^P necessitates frequent production of probes (5). As a result, we sought to implement and optimize a non-radioactive northern blotting method.

Alternative, non-radioactive approaches for probe-labeling (e.g. digoxigenin, biotin) and detection (e.g. colorimetric, fluorescence, luminescence) have been described (6-14). For our application, we selected labeling with biotin due to its strong and specific interaction with streptavidin and wide availability of biotin-streptavidin reagents (15). We chose luminescence detection due to its high sensitivity and prevalence in laboratory applications - the same method is widely used for western-blotting (16). Chemiluminescence detection can be fast and sensitive with high signal and low background.

In this report, we present an optimized non-radioactive northern blot method that utilizes chemiluminescence detection. This chemiluminescent northern approach (hereon referred to as Chemi-Northern) uses biotinylated antisense nucleic acid probes. These probes can be made “in house” or purchased from commercial sources. After hybridization to the target RNA on the blot, the probes are detected using streptavidin conjugated to horseradish peroxidase and enhanced chemiluminescence substrate. Importantly, all of the reagents and equipment are widely available and commonplace in molecular biology laboratories. To demonstrate the utility of Chemi-Northern, we test its dynamic range and sensitivity for multiple RNAs (including reporter gene and endogenous mRNAs and non-coding RNA) present in RNA isolated from human and *Drosophila* cells. Our results show that attomole sensitivity is possible with excellent linearity in response with detection times of only a few seconds. We show that Chemi-Northern accurately and reproducibly measures regulation of a reporter mRNA by the sequence-specific RNA-binding proteins PUM1 and PUM2. Furthermore, we use the method to measure enrichment of a reporter mRNA with the poly(A) binding protein, PABPC1. Our results show that Chemi-Northern is a versatile, effective method to analyze RNAs.

## Results

### Synthesis of biotinylated nucleic acid probes

Chemi-Northern detection uses biotinylated antisense nucleic acid probes - either RNA or DNA. Labeled probes of single-stranded RNA, transcribed *in vitro*, are commonly used for high sensitivity and ease of production (17-19). To generate an RNA probe, a DNA template for in vitro transcription is first produced with the appropriate promoter sequence. Suitable templates include PCR products, plasmids, or DNA oligonucleotides. We used the bacteriophage T7 RNA polymerase to transcribe probes for firefly luciferase (Fluc) and nano-luciferase (Nluc) reporter mRNAs from PCR-generated templates amplified with specific primers; each reverse primer contained a 5’ T7 promoter sequence 5’-TAATACGACTCACTATAGGG (the underlined G is the first nucleotide of the transcript)(see Methods). The Nluc and Fluc probes correspond to those that we previously used for radioactive northern blotting (20,21). The RNA probes were labeled with biotin during transcription using a mixture of NTPs with a biotin-16-UTP:UTP ratio of 1:2. After transcription, the RNA probes were purified using a spin column-based purification kit. Unlike radioactive probes, the biotinylated probes can be conveniently measured by UV absorbance (biotin does not absorb in the UV spectrum) and are stable when stored at -80 °C. The purified probes were also visualized by denaturing formaldehyde-MOPS agarose gel electrophoresis (**Figure. 1**), confirming their proper size and purity. Though biotin is randomly incorporated by T7 polymerase, the resulting probes run as a diffuse, single band of slightly higher apparent molecular weight, as previously observed (14).

As an alternative probe format, we purchased synthetic antisense DNA oligonucleotides with a 5’ biotin modification corresponding to β-actin (ACTB) mRNA and 7SL noncoding RNA. The general guidelines for DNA oligo probe design include lengths of 25-45 nucleotides with melting temperature (Tm) between 78-90 °C and GC content ranging from 45-65%, as recommended by the manufacturer of the hybridization buffer (see Methods). As such, the β-actin DNA probe was 41 nt with 59% GC content and a Tm of 71 °C. The 7SL DNA probe was 45 nt with a 60% GC content and Tm of 75 °C.

### Dynamic detection of reporter mRNAs expressed in cells using Chemi-Northern

To test the efficacy of the Chemi-Northern approach, we first sought to detect luciferase reporter mRNAs expressed in human and *Drosophila* cells. To assess the dynamic response of detection, the HCT116 colon cancer cells were transfected with increasing amounts of Nluc reporter expression plasmid (from 250 to 2000 ng of transfected plasmid per well of a 6-well plate). After 48 hours, total cellular RNA was purified using magnetic bead-based isolation, which incorporates on-bead DNase digestion to remove DNA. As a negative control, total RNA from untransfected cells was also analyzed. The concentration of the cellular RNA was determined by UV absorbance and the integrity of the RNA was assessed by ethidium bromide stained, denaturing formaldehyde-MOPS agarose gel electrophoresis of 5000 ng of each RNA sample. Imaging of the gel demonstrated that the 28S and 18S ribosomal RNAs (rRNAs) were intact (**Figure 2A and 2B**).

**Figure 2.**
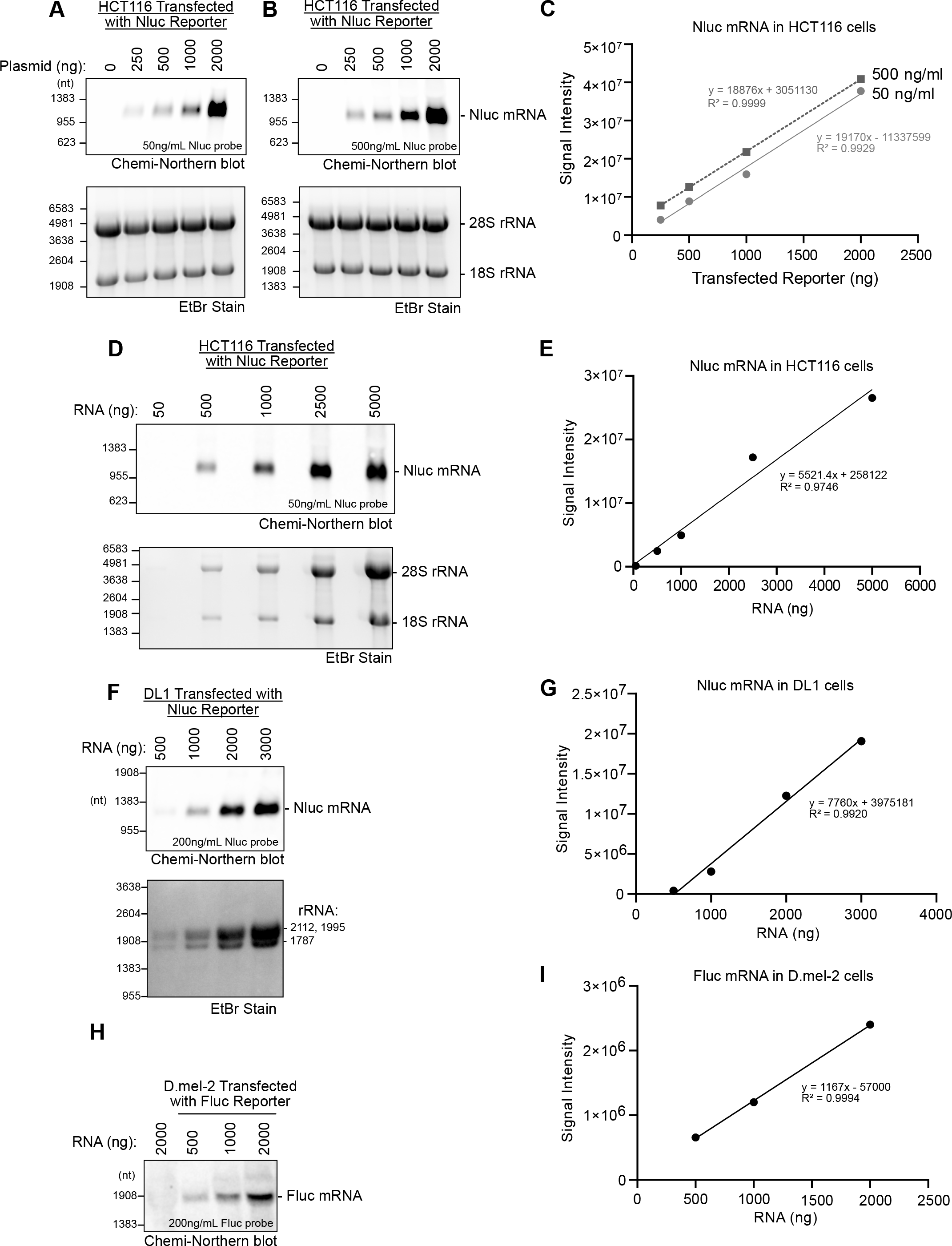
Dynamic response range of mRNA detection by Chemi-Northern using biotinylated RNA probes. HCT116 cells were transfected with increasing amounts (250, 500, 1000, and 2000 ng) of nano-luciferase reporter plasmid (Nluc), as indicated at the top. The ‘0’ ng condition indicates mock transfected cells with Nluc reporter. A total of 5 µg of purified cellular RNA was loaded into each lane of the denaturing formaldehyde-MOPS agarose gel and detected via Chemi-Northern using 50 ng/mL (A) or 500 ng/mL (B) of biotinylated antisense Nluc RNA probe. Chemi-Northern blot of expressed Nluc mRNA (639 nt +pA tail) at 1 sec exposure time is shown in the upper panels and ethidium bromide (EtBr) stain of ribosomal RNA (18S, 1869 nt and 28S rRNA, 5070 nt), serving as a loading control and indication of RNA integrity, are shown in the lower panels. (C) Graph showing the linear relationship between transfected reporter (x-axis) and signal intensity (y-axis) of Chemi-Northern blot at two different probe concentrations (50 and 500 ng/ml). Reported are the linear regression equation, y=mx+b and coefficient of determination, R^2^. All blots were quantified using AzureSpot Pro and graphs were created using GraphPad Prism software. (D) HCT116 cells were transfected with 1.25 µg of Nluc reporter plasmid and the purified total RNA was titrated (50, 500, 1000, 2500, and 5000 ng) and expressed Nluc mRNA was detected at 1 sec exposure by Chemi-Northern blot (upper panel) and EtBr stain (lower panel). (E) Quantitation and linear fit between total RNA mass (ng, x-axis) and signal intensity (y-axis). RNA was analyzed from *Drosophila* DL1 cells (F-G) or D.mel-2 cells (H-I) transfected with Nluc (806 nt +pA tail) or Fluc (1773 nt + pA tail) reporters using Chemi-Northern. Purified total cellular RNA was titrated as indicated at the top of the gels. Note that in panel F, the bottom panel shows the *Drosophila* rRNA species stained with EtBr, of which the 28S rRNA (3945 nt) is naturally processed into two fragments (1787 nt and 2112 nt) whereas the 18S rRNA is 1995 nt, as previously documented (30,31). (G, I) Quantitation of signal intensity relative to the amount of total cellular RNA analyzed.

The RNA was transferred from the gel to a positively charged nylon membrane by downward capillary blotting and crosslinked by exposure to UV (22). The membrane was then “pre-hybridized” by incubating with hybridization buffer. For hybridization, we initially tested two concentrations of biotinylated riboprobes (50 ng/mL (241 pM) and 500 ng/mL (2.41 nM)) to determine if there was an effect on the detection range. After probe hybridization, the membrane was extensively washed. To detect the mRNA-bound biotinylated probe, the membrane was blocked and then incubated with conjugated streptavidin-HRP followed by enhanced chemiluminescence substrate. The membrane was imaged using a CCD-camera based western blot imaging system. The resulting images were analyzed using software supplied with the imaging system, enabling quantitation of the signal intensity for each band and background correction.

For both 50 ng/ml (**Figure 2A**) and 500 ng/ml (**Figure 2B**) of probe, Nluc mRNA was detected within 1 second with nearly identical signal intensities and linear response ranges from 250 to 2000 ng of transfected reporter plasmid (**Figure 2C**). The Nluc mRNA signal was only present in transfected samples, demonstrating specificity, with high signal intensity and low background noise. The results also indicate that the probe is in excess under these conditions.

To further test Chemi-Northern detection, we analyzed a titration of total cellular RNA (50-5000 ng) from HCT116 cells transfected with 1.25 μg of Nluc expression plasmid. Nluc mRNA was detected by a 1 second exposure in as little as 250 ng of total RNA (**Figure 2D**) with signal intensity that was proportional to the amount of total (**Figure 2E**). We also used Chemi-Northern to detect Nluc (**Figure 2F**) and Fluc mRNAs (**Figure 2H**) expressed in cultured *Drosophila* cells. This experiment used a shorter 187 nt biotinylated RNA probe that hybridized to the Nluc 3′UTR, which was previously used for radioactive northern blotting (20). Quantitation of these blots provides additional evidence for the linear response of the assay (**Figures 2G and 2I**).

### Detection of endogenous RNAs using DNA oligonucleotide probes by Chemi-Northern

We used Chemi-Northern to detect two endogenous RNAs, including the β-actin mRNA and the 7SL noncoding RNA. For this application, we used biotinylated synthetic DNA oligonucleotide probes in a hybridization buffer optimized for oligonucleotide probes. β-actin mRNA and 7SL ncRNA were readily detected in as little as 500 ng of total HTC116 RNA using 5 nM of each probe (**Figure 3A**) with excellent linear response in signal intensity proportional to the amount of total RNA (**Figure 3B and 3C**). Similar results were obtained using a biotinylated DNA oligonucleotide to the *Drosophila* 7SL ncRNA (**Figure 3D and 3E**). Thus, Chemi-Northern effectively detects endogenous RNAs using synthetic probes with a single biotin label.

**Figure 3.**
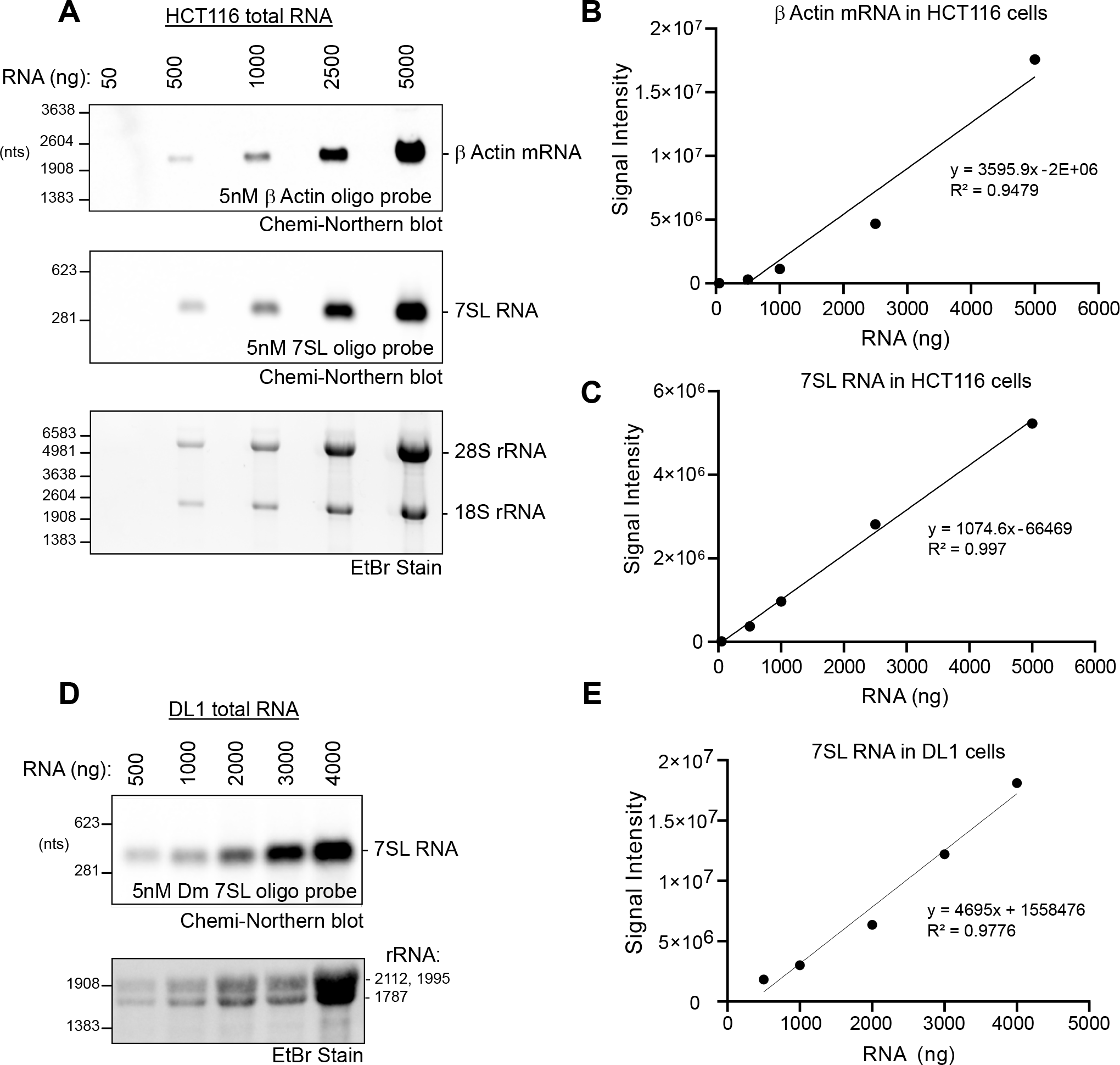
Chemi-Northern using biotinylated DNA probes detects endogenous messenger and non-coding RNAs. (A) Total RNA was purified from HCT116 cells and titrated (50, 100, 1000, 2500, and 5000 ng) onto a formaldehyde agarose gel followed by Chemi-Northern blotting using 5’ biotinylated DNA oligonucleotide antisense probes for endogenous β-actin (ACTB) mRNA at 85 sec exposure (1812nt +pA tail, upper panel) and 7SL ncRNA at 1 sec exposure (299 nt, middle panel). EtBr staining of ribosomal RNAs is shown in the lower panel. Quantitation and graphs showing linear fit of the total RNA mass (ng) versus signal intensity of the ACTB (B) and 7SL (C). (D) Total RNA of *Drosophila* DL1 cells titrated and endogenous 7SL (299 nt) and detected via Chemiluminescent Northern blot at 1 sec exposure. E) Quantitation of signal intensity relative to total cellular RNA.

### Attomole sensitivity of Chemi-Northern detection

We next wished to test the sensitivity range of Chemi-Northern detection. To do so, we transcribed and purified an RNA corresponding to Nluc mRNA **(Figure 4A)**. We then analyzed a titration of Nluc mRNA from 50-1690 attomoles (0.01-0.35 ng) (**Figure 4B**). The Chemi-Northern using the biotinylated Nluc antisense probe detected as little as 240 attomoles (0.05 ng) of RNA with a linear response observed across the range (**Figure 4B, C**).

**Figure 4.**
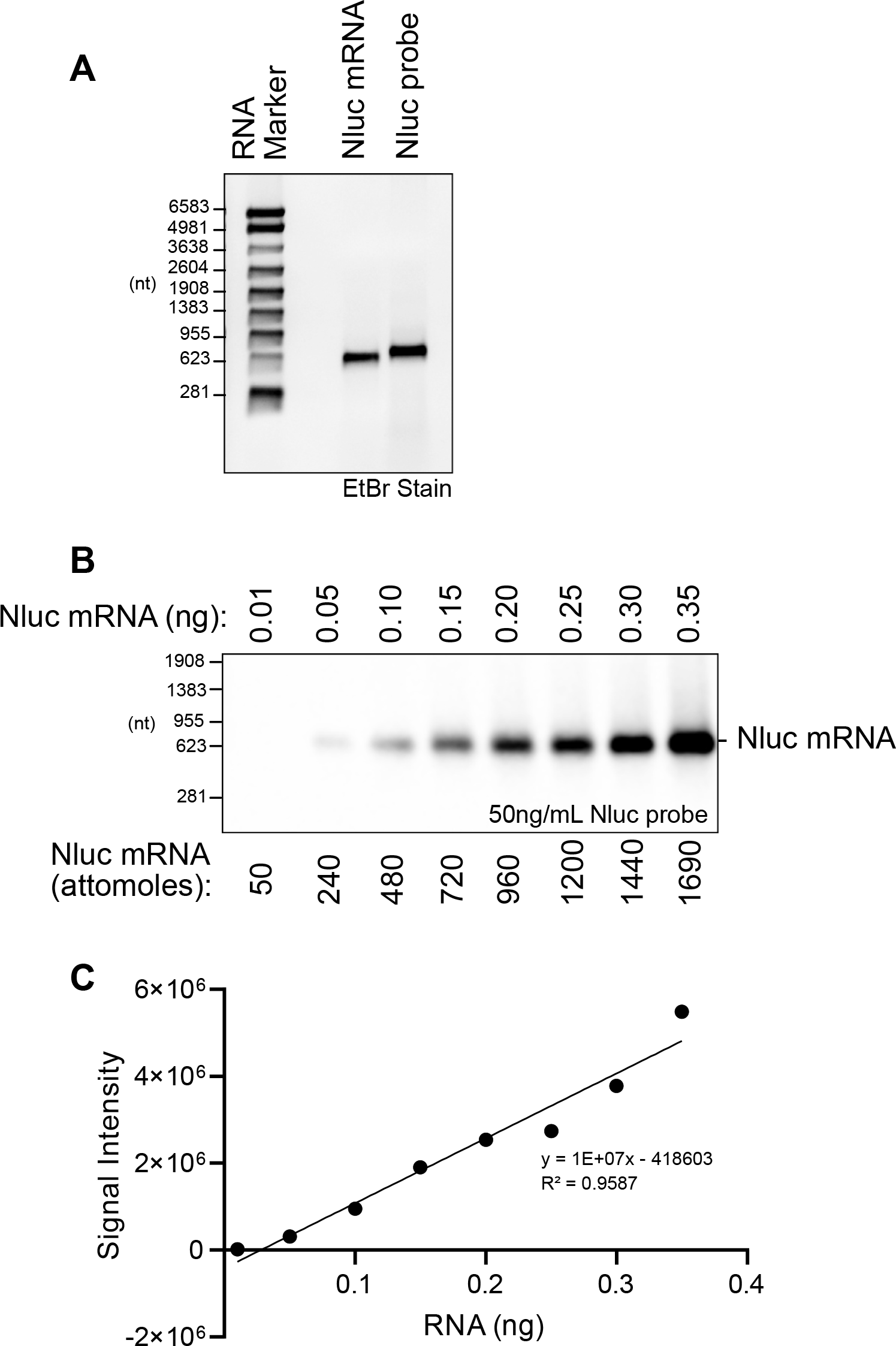
Attomole sensitivity of Chemi-Northern detection. (A) Ethidium bromide stained (EtBr) denaturing formaldehyde-MOPS agarose gel of 500 ng in vitro transcribed, purified synthetic Nluc mRNA and biotinylated Nluc antisense probe RNA (646 nt), confirmed proper size, purity, and quality. RNA size marker is included with nucleotide (nt) lengths indicated on the left. (B) Chemi-Northern detection of titrated synthetic Nluc mRNA at 6 sec exposure. The amount of the Nluc mRNA is indicated, with mass at the top and moles at the bottom. (C) Measured signal intensities in panel B are graphed relative to the amount of mRNA. Linear regression analysis was performed to assess the dynamic response, with the resulting line, equation, and coefficient of determination, R^2^.

We reasoned that the sensitivity of detection by Chemi-Northern could be enhanced by incorporating more biotinylated-uridine into the RNA probe. To test this idea, we transcribed a 197 nt Nluc 5ʹUTR RNA probe in reactions containing increasing ratio of biotinylated UTP to unlabeled UTP, while maintaining the same overall concentration. The probes were successfully produced at ratios of biotinylated UTP:UTP spanning 1:2 to 3:1, but not at 1:0 (**Figure 5A**). We noted a slight decrease in mobility of the probe produced at 3:1 ratio, indicative of the more densely incorporated bulky biotinylated uridine. We then probed identical blots with three amounts of total cellular RNA from HCT116 cells transfected with the Nluc reporter gene with either the 1:2 or 3:1 Nluc 5ʹUTR RNA probes. The same blots were probed with 7SL DNA oligo probe, serving as an internal control. The blots were visualized side-by-side under identical conditions, revealing stronger signal from the 3:1 Nluc 5ʹUTR RNA probe (**Figure 5B**). Quantitation of the signal intensities for each RNA amount, normalized to corresponding 7SL signal, demonstrates an average 5.4-fold increase in detection by the more heavily biotinylated 3:1 probe (**Figure 5C**).Thus, the sensitivity of Chemi-Northern can be enhanced by optimizing the degree of biotinylation of the RNA probes. We find that the 1:2 ratio works well for producing most probes. When increased sensitivity is needed, we recommend optimizing biotinylation of the probe to maximize label incorporation.

**Figure 5.**
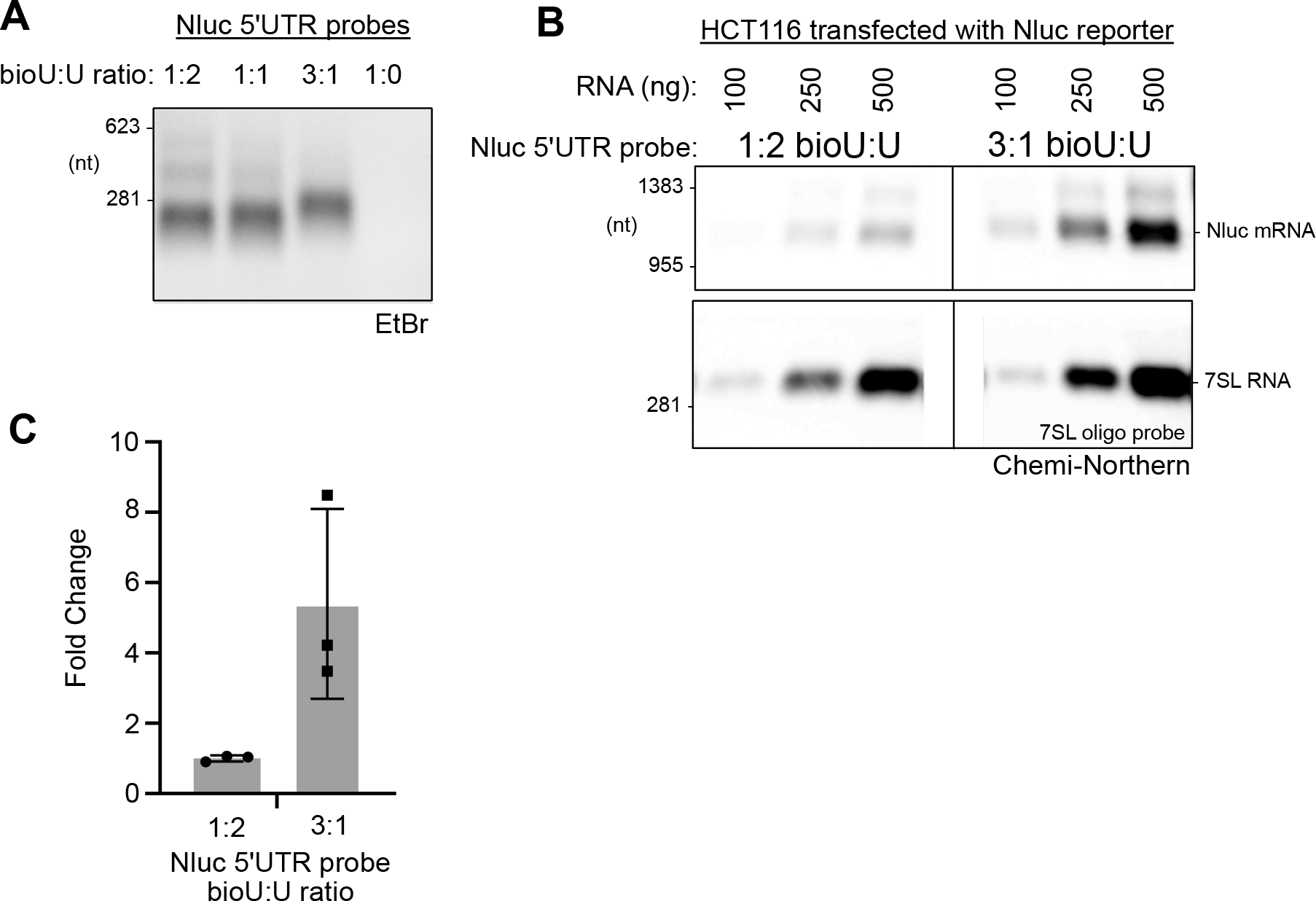
Enhanced sensitivity by optimizing biotin incorporation into RNA probe. (A) Ethidium bromided stained denaturing formaldehyde-MOPS agarose gel of Nluc 5ʹUTR RNA probes that were transcribed in vitro using the indicated ratios of biotinylated UTP (bioU) to unlabeled UTP. Total UTP concentration was the same in all reactions. (B) Chemi-Northern detection was performed on identical blots using Nluc 5ʹUTR probe with the indicated ratio of bioU:U. The gels were analyzed and imaged under identical conditions using the same exposure setting. 7SL RNA was detected using a biotinylated DNA oligo probe on the same blots, as a loading control. (C) Quantitation of the Chemi-Northern signals in panel B. Signal intensity for Nluc in each lane was normalized to its corresponding 7SL RNA signal. Fold change was then calculated relative to mean value for of the 1:2 bioU:U probe. Mean, SD, and the three data points for each condition are shown in the graph.

### Reproducible measurement of mRNA regulation and protein-RNA binding using Chemi-Northern

To demonstrate the utility of Chemi-Northern for analysis of mRNA regulation, and to assess the reproducibility of the results, we measured the effect of sequence specific RNA-binding proteins, human PUM1 and PUM2, on the levels of Nluc reporter mRNAs. PUM proteins are repressors that bind and degrade mRNAs that contain one or more Pumilio response elements (PRE: 5’-UGUANAUA) (21,23,24). We compared Nluc reporter mRNAs that contained 3 copies of the wild type PRE (Nluc 3xPRE) to a matched version wherein the first 3 nucleotides of the PRE were mutated to prevent PUM-binding (Nluc 3xPREmt, 5’-UGU changed to ACA). Consistent with our previous results (21,24,25), the Nluc 3xPRE mRNA was significantly and reproducibly reduced in three biological replicates relative to the mutant 3xPREmt version (**Figure 6A, B**).

**Figure 6.**
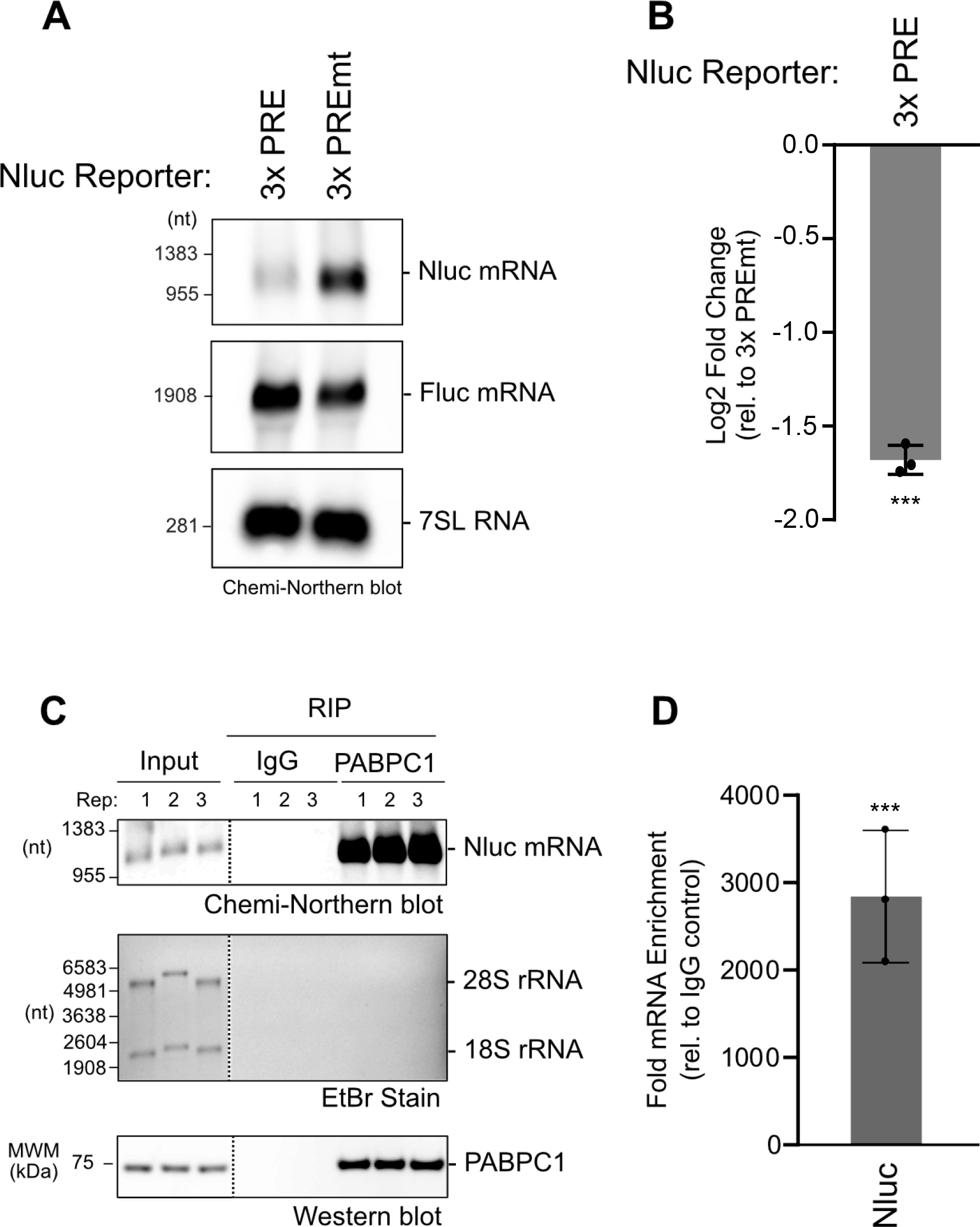
Chemi-Northern reproducibly measures mRNA regulation and binding by RNA-binding proteins. (A) Regulation of the Nluc reporter mRNA with three wild type Pumilio Response Elements (3xPRE) in its 3′UTR was compared to the mutant version using Chemi-Northern. The PREs are specifically recognized by endogenous sequence specific RNA binding proteins, PUM1 and PUM2, which repress by causing degradation of the mRNA, as previously documented (21,24,25,32). Reporters were expressed in HCT116 cells. The Fluc mRNA served as a control for transfection efficiency. Nluc mRNA was detected at 5 sec exposure and Fluc mRNA at 10 sec. The 7SL RNA served as an internal control for equivalent loading of the gel and was imaged at 5 sec exposure time. (B) Quantitation of Chemi-Northern in panel A and three additional biological replicates. Log2 fold change of the Nluc 3xPRE reporter was determined relative to the mutant version, Nluc 3xPREmt, in each sample was calculated as described in the Materials and Methods. Mean and standard deviation values are plotted, along with the values of each of the replicates. Statistical significance of p = 0.0005, based on paired two-tailed t test, is indicated by ‘***’. (C) Nluc mRNA is enriched by RNA coimmunoprecipitation (RIP) of human RNA-binding protein PABPC1 from HCT116 cells. The top image shows the Chemi-Northern detection of the poly-adenylated Nluc mRNA at 40 sec exposure in the input samples and the robust enrichment in the PABPC1 RIP samples, but not the IgG negative control RIPs. The middle image shows the ethidium bromide (EtBr) stained denaturing formaldehyde-MOPS agarose gel from input and RIP samples. Ribosomal RNAs are clearly visible in the inputs, but not RIP samples. The image at the bottom shows the western blot of PABPC1 protein, demonstrating its presence in the input samples and enrichment in the RIP samples. Dashed vertical lines in the panels indicate that the images were cropped to show relevant lanes. For each image, all of the lanes are from the same blot and exposure. (D) Quantitation of the Chemi-Northern in panel C demonstrates significant, reproducible detection of Nluc mRNA in the PABPC1 RIP samples. Nluc RNA signal intensity in each RIP sample was normalized to its corresponding input sample, then the fold enrichment in the PABPC1 RIP was calculated relative to the IgG negative control. Mean and standard deviation values are plotted, along with the values of each of the three biological replicates. Statistical significance of p = 0.0005, based on paired two-tailed t test, is indicated by ‘***’.

We also applied the Chemi-Northern to detect RNA-protein interactions in an RNA-coimmunoprecipitation (RIP) assay. For this experiment, we immunoprecipitated the poly-adenosine binding protein PABPC1, which interacts with the 3′ poly(A) tail of mRNAs to control their translation and stability. Cell extract was prepared from HCT116 cells that expressed poly-adenylated Nluc mRNA. PABPC1 was immunoprecipitated from the cell extract using a specific antibody and then the associated RNA was purified for the RIP and input materials. As a negative control, a mock RIP was performed from the cell extract using nonspecific immunoglobulin (IgG). Chemi-Northern detected Nluc mRNA in the PABPC1 RIP (**Figure 6C**), whereas ribosomal RNAs were not enriched (as detected by ethidium bromide staining of the gel). Western blot analysis confirmed the presence of PABPC1 in the RIP samples. Quantitation of the Chemi-Northern shows substantial and reproducible enrichment of the Nluc mRNA relative to the negative control (**Figure 6D**).

## Discussion

Northern blotting has proven to be a valuable methodology to detect and quantitate RNAs in many applications. The Chemi-Northern approach described here expands its usefulness. As examples, we demonstrate its versatility by analyzing intact RNA molecules - both mRNAs and ncRNAs, natural and engineered - in total RNA or RNA coimmunoprecipitation assays. Chemi-Northern has multiple advantages when compared to traditional Northern blotting and alternative RNA analysis methods. 1) The challenges associated with use of radioactive nucleotides are relinquished. 2) The procedure is simple. 3) The necessary reagents and equipment are minimal, widely available, and prevalent in molecular biology laboratories. 4) The cost per assay is relatively small compared to other techniques. Based on prices at the time of this publication, we estimate ∼$100 per blot for supplies. 5) Chemi-Northern probes are easy to create and are stable. 6) Chemiluminescence detection is fast, with exposure times as short as 1 sec. Overall, Chemi-Northern procedure can be completed in 2 days. 7) Chemi-Northern is sensitive: detection of attomole amounts of RNA is readily achievable. For lower abundance RNAs, detection can be boosted by optimizing biotin incorporation into the probe. We also suggest that enrichment of the target RNA, such as poly(A) selection of mRNAs (26), could enhance detection. 8) Our results show excellent linearity and dynamic range. The signal intensity is high with low background noise. Moreover, the measurements proved to be reproducible, enabling us to quantitate mRNA regulation by human PUM proteins and RNA-binding by PABPC1. We anticipate that the Chemi-Northern approach will be useful to the scientific community, empowering their research on RNA biology, gene regulation, and disease mechanisms.

### Experimental Procedures

#### Plasmids

The following plasmids were used in this study:

pNLP Nluc 3xPRE KM054 Enwerem et al, 2021 (21)

pNLP Nluc 3xPREmt KM055 Enwerem et al, 2021 (21)

pGL 4.13 Fluc KM052 Promega

pF5A empty vector ACG1092 Enwerem et al, 2021 (21)

pNLP MCS JAB1 Enwerem et al, 2021 (21)

pAC5.1 Fluc2 CAW023 Weidmann et al, 2012 (27)

pAC5.4 Nluc2 MCS RMA0056 Arvola et al, 2020 (20)

pIZ V5 H6 Themo Fisher Scientific

#### In vitro transcription of RNAs

DNA templates used to transcribe the biotinylated antisense RNA probes corresponding to Nluc and Fluc were generated by PCR using the nano-luciferase (Nluc) plasmid, pNLP MCS and Firefly luciferase (Fluc) plasmid, pGL4.13, respectively. The following primers were used for PCR, and the reverse primer incorporated the promoter for T7 RNA polymerase (underlined):

Nluc probe F: 5′-CACTCGAAGATTTCGTTGGGGAC

Nluc probe R: 5′-GGATCC<u>TAATACGACTCACTATAGGG</u>GATGCGAGCTGAAGCACAAGC

Fluc probe F: 5′-CGAGATGAGCGTTCGGCTGGCAGAA

Fluc probe R: 5′-GGATCC<u>TAATACGACTCACTATAGGG</u>CCGAAGCCGTGGTGAAATGGCA

Transcription from the Nluc template produces a 646 nt RNA probe that is complementary to the Nluc coding sequence. The Fluc template produces a 599 nt RNA probe complementary to the Fluc coding sequence.

The DNA template for transcription of the synthetic Nluc mRNA was generated using the following primers:

Nluc Sense F: 5′-GGATCC<u>TAATACGACTCACTATAGGG</u>AGACACTCGAAGATTTCGTTGGGGAC Nluc Sense R: 5′-GATGCGAGCTGAAGCACAAGC

Transcription from this template produces a 646 nt RNA within the Nluc coding sequence.

The DNA template for transcription of Nluc 5ʹUTR RNA probe was generated using the following primers:

Nluc 5ʹUTR probe F: 5′-CTCTGAGCTATTCCAGAAGTAGTGAGGAG

Nluc 5ʹUTR probe R: 5′-GGATCC<u>TAATACGACTCACTATAGGG</u>AGTACTCTAGCCTTAAGAGC

Transcription from this template generated a 197 nt probe.

For luciferase reporters in *Drosophila* cells, DNA templates were similarly used to transcribe biotinylated antisense RNA probes. The template for the Nluc 3′UTR probe was generated by PCR from the pAC5.1 Nluc MCS plasmid as previously described (20) using the following primers:

Nluc 3′UTR probe F: 5′-GGTTGAAGAGCAAGCCGC

Nluc 3′UTR probe R: 5′-GGATCC<u>TAATACGACTCACTATAGGG</u>GCGGCCAGCGGC

Transcription from this template generates an 187 nt probe with 124 nt complementarity to the 3′UTR of the Nluc reporter.

The template for the Fluc probe was generated by PCR from pAC5.1 Fluc pA plasmid using the following primers:

Fluc probe F: 5’-TAAGACACTGGGTGTGAACCAGCGCG

Fluc probe R: 5’-GGATCC<u>TAATACGACTCACTATAGGG</u>CTTGGCCTTAATGAGAATCTCGCGGA

Transcription from this template generates a 493 nt probe complementary to the Fluc coding region.

PCR generated DNA templates were purified using DNA clean and concentrator 25 spin columns (Zymo, D4065). The proper size of the PCR products were confirmed by agarose gel electrophoresis. The riboprobes were then transcribed using the MAXIscript T7 Transcription Kit (Thermo Fisher Scientific, AM1314). For a 20 µL transcription reaction, 1 μg of PCR template DNA was incubated with 2 µL Biotin RNA Labeling Mix (containing 10 mM each ATP, CTP, GTP, 6.5 mM UTP, and 3.5 mM Biotin-16-UTP, pH 7.5)(Roche, 11685597910), 2 µL 10x transcription buffer, and 2 µL T7 enzyme (30 U). Note that for **Figure 5**, the amounts of Biotin-16-UTP and UTP used were varied as indicated in the figure, while keeping the total UTP concentration at 10 mM. The reaction was incubated at 37 °C for 15 mins. The template DNA was degraded by adding 1 µL (2 U) Turbo DNase (Thermo Fisher Scientific, AM2238) with incubation at 37 °C for 10 mins.

The RNA was then purified using the RNA Clean & Concentrator-25 kit (Zymo, R1017) following the manufacturer’s instructions and eluted in 50 µL. The concentration of the RNA was measured by UV absorbance with a NanoDrop spectrophotometer (Thermo Fisher Scientific) and RNA integrity was visualized by electrophoresis and ethidium staining of RNA separated on a denaturing MOPS formaldehyde agarose gel (described below).

#### Cell Culture and Transfection

Human HCT116 cells (ATCC, CCL-247) were cultured in McCoy’s 5A modified medium (Thermo Fisher Scientific, 16600082) supplemented with 10% (v/v) fetal bovine serum (Genesee Scientific, 25-514) and antibiotics (100 U/mL penicillin and 100 µg/mL streptomycin, Thermo Fisher Scientific), at 37 °C with 5% CO_2_. HCT116 cells were seeded at 200,000 cells in 2 ml per well in a 6-well plate (USA Scientific, CC7682-7506) and, after 24 hours, were transfected with the amount of Nluc plasmid indicated in the figures. Cells were transfected using Fugene HD (Promega, E2312), following the method specified by the manufacturer and a ratio of 4 μl of Fugene HD to 1 μg of plasmid DNA. We note that for assays with varying amounts of transfected reporter plasmid, the total amount of transfected DNA was held constant by balancing with a control plasmid, pF5A empty vector. Unless otherwise noted, standard transfection conditions used 3μg total plasmid DNA containing 0.5 μg Fluc plasmid (pGL 4.13), 1.25 μg of Nluc plasmid, and 1.25 μg of pF5A empty vector. The transfected cells were grown for another 48 hrs and then RNA was purified for the Chemi-Northern analysis.

*Drosophila* D.mel-2 cells (Invitrogen) were cultured in Sf900III (Thermo Fisher Scientific, 12658-019) supplemented with antibiotics (25 U/mL penicillin and 25 μg/mL streptomycin)(Thermo Fisher Scientific) at 25 °C. D.mel-2 cells (2x10^6^ in 2 mL per well) were seeded in a 6-well plate and transfected with 5 ng pAC5.1 Fluc pA plasmid. Transfection master mixes contained 3 μg total plasmid DNA, Sf900III to 144 μL, and 2 μL of Fugene HD per 1 μg of total plasmid DNA. For experiments wherein the reporter plasmid was titrated, the total amount of transfected DNA was balanced using control plasmid, pIZ V5 H6 empty vector. The transfected cells were grown for an additional 48 hrs then RNA was purified for Chemi-Northern analysis.

*Drosophila* DL1 cells (also known as S1) were transfected as described above with the following alterations: cells were cultured in Schneider’s Drosophila Media (SDM, Thermo Fisher Scientific, 21720024) supplemented with 1x antibiotic-antimycotic (Thermo Fisher Scientific, 15240062), 1% (v/v) Glutamax (Fisher, 35-050-061), and 10% (v/v) heat-inactivated fetal bovine serum (Genesee, 25-514). Transfections contained 5 ng pAC5.4 Nluc pA plasmid, 5 ng pAC5.1 Fluc pA plasmid, pIZ V5 H6 control plasmid (up to 3 μg total DNA), serum-free SDM media, and 2 μL of Fugene HD per 1 μg of total plasmid DNA.

#### Purification of cellular RNA

HCT116 cells were harvested by washing twice with phosphate buffered saline (PBS) pH 7.4 (Thermo Fisher Scientific), followed by trypsinization with TrypLE (Thermo Fisher Scientific) for 5 mins, and centrifugation at 500 g for 5 mins. RNA was purified from the cell pellet using the SimplyRNA cells kit (Promega, AS1390) and Maxwell RSC instrument following the manufacturer’s protocol. For the on-bead DNase digestion, the amount of DNase I (Promega, Z3585) was doubled (10 μL total) to ensure removal of genomic and plasmid DNA. Purified RNA was eluted in 40 μL of nuclease-free water and quantified using a NanoDrop spectrophotometer.

#### Denaturing gel electrophoresis of RNA

RNAs were analyzed by denaturing gel electrophoresis in a chemical fume hood using a 1% (w/v) agarose gel (15 cm x 10 cm) containing 1x MOPS buffer (20 mM 3-(N-Morpholino)propanesulfonic acid hemisodium salt (MOPS) pH 7.0, 5 mM sodium acetate, and 1 mM EDTA) with 1.48% formaldehyde. Typically, 2.5 µg of total cellular RNA per sample was prepared in a final volume of 24 µL containing final concentrations of northern RNA sample buffer (0.04 μg/μL ethidium bromide, 23% formamide, 3% formaldehyde, 4.6 mM MOPS, 1.1 mM sodium acetate, and 0.2 mM EDTA), and northern RNA loading dye (2.1% glycerol, 4.2 mM EDTA and 0.01% (w/v) of Bromophenol Blue and Xylene Cyanol). Note that ethidium bromide was not added to the loading buffer for low amounts of RNA (<200 ng), such as RIP samples, because it can affect the migration of the RNA. Instead, those gels can be stained by soaking with ethidium bromide after electrophoresis. For higher amounts of RNA, ethidium bromide did not appreciably impede electrophoresis, transfer, or hybridization, as previously reported (28). RNA molecular weight markers (Promega, G3191) were included. The RNA samples were heated at 70 °C for 10 mins before loading the entire volume on the agarose gel. The RNA was separated by gel electrophoresis in 1x MOPS running buffer at 95 V for 1.5 hrs. The size, integrity, and equivalent loading was assessed by ethidium bromide staining and UV visualization of the ribosomal RNA.

#### Blotting of RNA to membrane

After gel electrophoresis, the RNA was transferred from gel to positively charged nylon membrane (Immobilon-Ny+, Millipore, INYC00010) by downward capillary transfer overnight in 20x SSC buffer containing 3 M NaCl and 300 mM sodium citrate (pH 7.0) (22,29). Briefly, the northern blot transfer apparatus was assembled with a ∼1” high stack of cellulose chromatography paper (Thermo Fisher Scientific, 05-714-4). Two sheets of cellulose chromatography paper were soaked in 20x SSC and added to the top of the stack. The nylon membrane was then soaked in ddH_2_O for 5 mins, followed by 20x SSC for 5 mins, and then assembled on the top of the stack. The formaldehyde-agarose gel was carefully placed on top of the membrane using a roller to eliminate any bubbles. Two additional sheets of chromatography paper, soaked in 20x SSC, were added to the top of the gel. A wick of chromatography paper was placed on top with each end submerged in a container holding excess 20x SSC. After transfer, the nylon membrane (i.e. blot) was exposed to UV (120 J/cm^2^, λ= 254 nm) in a CL-1000 crosslinker (UVP) to crosslink the RNA to the membrane. To detect the RNA marker, its lane was removed from blot and visualized by staining with 0.25% w/v methylene blue in ddH_2_O for 5 mins, followed by washing in ddH_2_O until marker bands were clearly visible (10 mins).

#### Probe Hybridization

The RNA probes were described above. The following DNA oligonucleotide probes were also tested.

β-Actin biotinylated DNA oligo probe for human ACTB mRNA

5′ biotin-GATGGGGTACTTCAGGGTGAGGATGCCTCTCTTGCTCTGGG

7SL biotin DNA oligo probe for human 7SL ncRNA

5′ biotin-CTTAGTGCGGACACCCGATCGGCATAGCGCACTACAGCCCAGAAC

Dmel 7SL biotin DNA oligo probe for *Drosophila* 7SL ncRNA

5′ biotin-GATTGTGGTCCAACCATATCGGTTGGGCTGATAACGGCAGTCCAC

The blot was transferred to a hybridization bottle and incubated with rotation in 10 mL pre-warmed Ultrahyb Ultrasensitive hybridization buffer (Thermo Fisher Scientific, AM8670) at 68 °C for 1 hr for RNA-based probes or 10 mL of pre-warmed Ultrahyb oligo hybridization buffer (Thermo Fisher Scientific, AM8663) at 42 °C for 1 hr for DNA oligo probes. RNA based probes were typically hybridized at 50 ng/ml, unless otherwise noted in the figures. For DNA oligo based probes, 5 nM was used. Hybridization proceeded for 12 or more hours at 68 °C (RNA probe) or 42 °C (DNA oligo probe) in a rotisserie hybridization oven (Thermo Fisher Scientific).

#### Chemiluminescent Detection

After probe hybridization, the buffer was removed from the bottle and then the blot was washed twice with 2xSSC and 0.1% SDS and then twice with 0.1xSSC and 0.1% SDS. Each wash was performed at 68 °C (RNA probe) or 42 °C (DNA oligo probe) using 25 mL of the stated buffer for 15 minutes each in a rotisserie hybridization oven. The Chemiluminescent Nucleic Acid Detection Module (Thermo Fisher Scientific, 89880) was used to detect the biotinylated probe on the blot with streptavidin-HRP and enhanced chemiluminescence using the manufacturer’s instructions and supplied reagents. Detection was performed at room temperature. The blot (typically 10x10 cm) was first incubated with 16 mLs of blocking buffer in a clean tray for 15 mins at room temperature with gentle rocking. Next, 50 μl of streptavidin-HRP was added to the tray containing the blot and blocking buffer and was incubated with rocking for 15 mins. The blot was then washed once with 20 mL of the supplied 1x wash buffer for 5 minutes with gentle rocking. The blot was then transferred to a new clean tray and washed four more times. The blot was transferred to a new tray containing 20 mL of the supplied equilibration buffer and incubated for 5 mins with rocking. During the equilibration, 8 mLs of substrate working solution per blot was prepared in a conical tube with equal volumes of the supplied chemi luminol/enhancer solution and stable peroxide solution. The substrate mixture was pipetted into a new tray using the puddle method with the blot placed face down into the liquid for 5 mins without rocking. The blot was wrapped in plastic wrap, placed onto a glass plate, and visualized using a chemiluminescence imaging system (Azure 300 Chemiluminescent Imager, Azure Biosystems) starting at 1 sec exposure for RNA probes and 10 sec exposure for oligo probes. Exposure times were adjusted as needed.

#### RNA Immunoprecipitation

HCT116 cells were seeded in a 6-well plate and pNLP MCS Nluc reporter plasmid transfected as described above. Two wells were harvested for each RNA coimmunoprecipitation (RIP) analysis. For each sample, 8.3 µl Dynabeads protein A (Thermo Fisher Scientific, 10001D) were washed 3 times with 250 µl 1xPBS + 0.1% Tween-20 and incubated in 100 µl 1xPBS + 0.1% Tween-20 + 2 µg PABPC1 antibody overnight (Abcam, ab6125), rotating at 4°C. The next day the HCT116 cells were collected by centrifugation and lysed by cell disruption in 400 µl of lysis buffer (50 mM Tris-HCl pH 8.0, 500 mM NaCl, 0.5% Triton-X100, 1 mM EDTA, RNase Inhibitor (RNasin, Promega) with 2x complete protease inhibitor cocktail (Roche, 11836153001). Cell debris was removed by centrifugation at 10,000 g for 10 mins and the supernatant was then passed through a 0.45-micron filter at 4,000 g for 3 mins. The protein concentration of the extract was measured using the DC-Lowry assay (BioRad). For each sample, 10 µl of lysate was set aside for the input sample. Next, 300 µg of total protein in 200 µl of lysis buffer was added to the protein A beads and incubated for 2 hrs with end over end rotation at 4°C. The supernatant was removed from the RIP samples and the beads were subsequently washed 6x with 500 µl of lysis buffer without RNasin. Washed beads were suspended in 20 µl, 5 µl of which was reserved for western blot analysis (described below). RNA was purified from the input sample and the remaining beads using the RNA clean and concentrator 5 kit (Zymo, R1013) following the manufacturer’s instructions, including the on-bead DNase I digestion. RNA was eluted in a final volume of 10 µl. The entire RIP RNA sample was used for northern blotting, along with 1% of the RNA purified from the input sample. Ethidium bromide was not included in the sample loading buffer for gel electrophoresis of the RIP samples.

#### Analysis and quantitation of chemi-northern blots

The chemi-northern blots were analyzed using AzureSpot Pro analysis software (Azure Biosystems). Individual lanes were identified automatically and adjusted manually as needed. Band signals were detected using the fixed width set to 50. The background was corrected using the rolling ball method with the radius set to 5%. The background corrected signal intensities (volumes, measured in arbitrary units) from titration experiments were plotted and used to determine the linearity of signal response. For northern blot quantitation of PUM repressive activity, the Nluc and Fluc signal intensities were normalized to their respective loading controls, 7SL ncRNA, on the corresponding blots. For each sample, the normalized Nluc reporter signal was then divided by the normalized Fluc signal, to correct for variation in transfection efficiency. The resulting relative response ratios were then used to calculate the log2 fold change in reporter Nluc activity between the PUM-regulated Nluc 3xPRE reporter and the unregulated PRE mutant reporter, Nluc 3xPREmt. Three biological replicates were performed for each measurement.

For the PABPC1 RIP analysis, the Nluc mRNA signal intensities of the IgG negative controls and RIP samples were normalized to their respective inputs. The fold enrichment of the normalized PABPC1 RIPs relative to the normalized IgG controls was then determined. Three biological replicates were performed for each measurement.

### Western blot analysis

For western blotting of RIP samples, proteins were separated by SDS-PAGE on a 4-20% gradient polyacrylamide gel (Bio-Rad) and transferred to a PVDF membrane (Immobilon P, Millipore) at 60V constant for 1.5 hrs at 4 °C. The membrane was then blocked for 1 hr in in 5% w/v nonfat powdered dry milk, 1xPBS, and 0.1% Tween-20 (PBST) and incubated in PABPC1 primary antibody (Abcam, ab6125) diluted to 1:1,000 in PBST overnight at 4 °C. The membrane was washed 3 times 10 mins with PBST and incubated with Anti-Mouse Ig HRP secondary (Mouse Trueblot, Rockland) diluted 1:10,000 in PBST. The membrane was washed again 3 times 10 mins with PBST and visualized by enhanced chemiluminescence using Immobilon Western Chemiluminescent HRP Substrate (Millipore).

## Supporting Information

This article does not contain supporting information.

## Acknowledgements

We thank Drs. Eric Wagner and Rene Arvola for their feedback and critiques on this manuscript prior to publication.

## Author Contributions

K.M. conceptualization, investigation, methodology, visualization, formal analysis, writing, review, editing

R.C. conceptualization, investigation, methodology, visualization, formal analysis, writing, review, editing

E.D. investigation, methodology, visualization, formal analysis, review, editing

A.G. funding acquisition, project administration, supervision, conceptualization, visualization, writing, review, editing

## Funding and additional information

This research was supported by National Institutes of Health grants 1R01GM145835 and 1R01GM144466 to A.C.G. The content is solely the responsibility of the authors and does not necessarily represent the official views of the National Institutes of Health. E.B.D was supported by University of Minnesota’s Targets of Cancer Training Program, NIH grant T32 CA009138.

